# Network plasticity in the honey bee antennal lobe during odor presentation

**DOI:** 10.1101/2022.11.17.516890

**Authors:** Marco Paoli, Yuri Antonacci, Luca Faes, Albrecht Haase

**Author notes:** **Correspondence:** Albrecht Haase, Marco Paoli.

## Abstract

Odorant processing presents multiple parallels across animal species, and insects became relevant models for the study of olfactory coding because of the tractability of the underlying neural circuits. Within the insect brain, odorants are received by olfactory sensory neurons and processed by the antennal lobe network. Such a network comprises multiple nodes, named glomeruli, that receive sensory information and are interconnected by local interneurons participating in shaping the neural representation of an odorant. The study of functional connectivity between the nodes of a sensory network *in vivo* is a challenging task that requires simultaneous recording from multiple nodes at high temporal resolutions. Here, we used fast two-photon microscopy to follow the calcium dynamics of antennal lobe glomeruli and applied Granger causality analysis to assess the functional directed connectivity among network nodes during and after olfactory stimulation. Our findings show that there is a significant causal coupling between network nodes also in the absence of olfactory stimulation. However, connectivity patterns change upon odorant arrival, showing an increase in connection density and conveying odorant-specific information. Furthermore, we show that causal functional connectivity is unique in individual subjects, and cannot be detected upon artificial glomerular scrambling across individuals. This demonstrates that network states do not consist solely of correlations of mean glomerular responses, which are known to be conserved across bees.

## Introduction

Honey bees are eusocial insects strongly relying on olfaction, *e*.*g*. for foraging, social communication, or mating. Their olfactory system has been extensively studied in the past decades, and it is structured along different anatomical and functional layers of odor processing (Paoli and Galizia, 2021). Volatile chemicals are received by olfactory receptors located on the antennae, where they are received by ∼60,000 olfactory receptor neurons (ORNs). Then, the olfactory input is forwarded to the first processing center, the antennal lobe (AL), where ORNs bearing the same olfactory receptor converge into one of about 160 glomeruli, the functional nodes of the AL processing network. Incoming neuronal inputs are further processed by ∼4,000 local neurons (LNs), and the resulting signal is forwarded by ∼800 projection neurons (PNs) into higher-order brain centers like the mushroom bodies (MBs), where information from different sensing modalities is integrated and long-term memory is formed, and the lateral horns (LHs), most likely involved in odor valence coding.

The development of *in vivo* calcium imaging in the honey bee (Galizia et al., 1999b; Paoli et al., 2017) allowed simultaneous recording of the activity of multiple glomeruli, revealing the basic principles of olfactory coding, such as the stereotypical odorant-specific glomerular response maps (Galizia et al., 1999b), corresponding to the time-averaged PN firing rates (Moreaux and Laurent, 2007), the representation of information on the odors’ chemical properties (Sachse et al., 1999) and concentration (Sachse and Galizia, 2003), or the link between similarity of perception and neural representation of an olfactory cue (Guerrieri et al., 2005). With the implementation of fast scanning two-photon imaging, the temporal resolution has been increased by more than an order of magnitude, passing from ∼6 up to 100 Hz (Haase et al., 2011), thus providing the access to fine temporal properties of the olfactory code. This allowed identifying the contribution of glomerular response latencies to the odor code (Paoli et al., 2018), and to access the time-frequency domain of the glomerular responses, which revealed stereotypical odorant-specific time-frequency response patterns (Paoli et al., 2016b). In addition, fast calcium imaging allows exploring AL network dynamics so far precluded to more traditional approaches: although electrophysiology provides very high temporal resolution, it allows recording from only a limited number of network nodes at once; conversely, wide-field calcium imaging grants simultaneous access to multiple nodes with sub-cellular spatial resolution, but the limited temporal resolution makes time-resolved correlation measurements and causality analysis impossible.

Previous attempts to investigate intra-glomerular connectivity relied on a modeling approach, *e*.*g*. matching ORNs and PNs signals to investigate the optimal arrangement of the local inhibitory network (Linster et al., 2005), or by combining local stimulation/inhibition and global recording to assess connectivity properties of the local network (Girardin et al., 2012). Here, we took advantage of the high spatial and temporal resolution of two-photon calcium imaging and identified the network of functional glomerular coupling based on Granger causality (GC) analysis. Granger causality was computed between pairs of glomeruli for each individual and across multiple odorants. This allowed studying connectivity patterns in the AL network *in vivo*, and their dynamics during, and after an olfactory stimulation. Finally, we could assess the stimulus information content of the elicited connectivity network across odorants and subjects.

## Material and methods

### Bee preparation

Forager honey bees *Apis mellifera* were collected at the outdoor hives of the institute (CIMeC, Rovereto, Italy) on the day of the experiment. They were cold-anesthetized at −20°C, placed on custom-made Plexiglass holders, and their heads were fixed with soft dental wax to avoid movements (Paoli et al., 2017). After opening a small window in the head cuticle, glands and tracheas were gently displaced to expose the bee brain, and the tip of a pulled glass capillary coated with Fura-2-dextran 10 kDa (ThermoFisher Scientific) was inserted between the medial and lateral mushroom body calyces to label the projection neurons tracts (Sachse and Galizia, 2002). Then, the head capsule was closed and sealed with *n*-eicosane (Sigma-Aldrich) to prevent brain desiccation. Bees were fed with 20 μL of a 50% sugar/water solution and kept overnight at room temperature in a dark and humid environment. On the following day, the window cuticle was re-opened and the antennal lobe ipsilateral to the injection site was exposed for imaging. The brain was covered in transparent silicon (Kwik-Sil, WPI) to prevent brain movements and desiccation. Note that this procedure has an intrinsic variability that depends on the amount of Fura-2 delivered by the experimenter and on the number of labeled projection neurons. Still, multiple studies have shown convergent results on odor-elicited glomerular responses, indicating that such a procedure remains valid to investigate olfactory processing in the honey bee antennal lobe (Hourcade et al., 2009; Locatelli et al., 2013; Paoli et al., 2016a; Bestea et al., 2022).

### Olfactory stimulation

Olfactory stimuli were delivered with a custom-built olfactometer comprising eight odor channels and controlled via a custom-made LabVIEW (National Instruments) interface, which synchronized stimulation protocol with the imaging acquisition (Paoli et al., 2017). For this study, we used six ecologically relevant odorants known to elicit distinct responses among the antennal lobe glomeruli: 1-hexanol (1HEX), 3-hexanol (3HEX), 1-nonanol (1NON), isoamyl acetate (ISOA), acetophenone (ACTP), and benzaldehyde (BZDA) (all from Sigma-Aldrich) (Galizia et al., 1999b; Paoli et al., 2018). Also, they provide different degrees of structural and functional variability: 1-hexanol and 3-hexanol have the same carbon length but vary for the position of the hydroxyl group, 1-hexanol and 1-nonanol are primary alcohols with different chain lengths; acetophenone and benzaldehyde both have a benzene ring, but with a ketone and an aldehyde group, respectively; isoamyl acetate is the major component of the alarm pheromone (Boch et al., 1962). Odorants were presented in a 1 s ON / 9 s OFF protocol and each odorant was delivered for 30 consecutive trials. All odorants were diluted 1:200 in mineral oil (Sigma-Aldrich). These concentrations were chosen to be well above the receptor sensitivity threshold and below the saturation level (Sachse and Galizia, 2003).

### Calcium imaging acquisition and data processing

Optical imaging was performed via a two-photon imaging platform based on an Ultima IV microscope (Brucker). The calcium sensor Fura-2 was excited at 800 nm and fluorescent changes were recorded with a photomultiplier (Hamamatsu) at 525 ± 20 nm. Line scanning was performed along a user-drawn line crossing all glomeruli in the imaging field and scanned at a rate of 100 Hz. Individual glomeruli were identified and labeled according to the honey bee reference atlas (Galizia et al., 1999a). To allow for cross-subject comparisons, the analysis was limited to the 10 glomeruli that could be identified, with a few exceptions, in all imaged bees. For the analysis, a whole imaging session, comprising 30 repetitions of 6 odorants, was decomposed into individual stimulation trials. The average fluorescence signal from each identified glomerulus and for each stimulation trial was normalized with respect to the mean pre-stimulus baseline (Δ*F*/*F*; the interval of 1 s before stimulus onset was used to define the pre-stimulus baseline). Data are always presented as the negative relative fluorescence change (-Δ*F*/*F*) so that excitatory responses correspond to positive changes and inhibitory responses to negative ones.

### Measuring Granger causality

The calcium time series recorded from *M* =10 glomeruli for each animal were described as a stationary vector stochastic process *Y*_*n*_ = [*Y*_1,*n*_, …, *Y*_*M,n*_]^*T*^, where *n* denotes the time stamp. The temporal statistical structure of this process was described using the vector autoregressive (VAR) model:

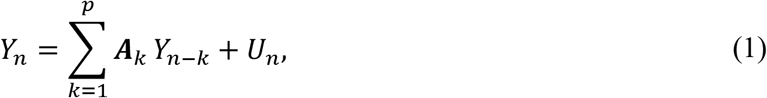

where *A*_*k*_ is an *M* × *M* matrix containing the coefficients that relate the present state of the process, *Y*_*n*_, to its past state evaluated at lag *k, Y*_*n*−*k*_, and *U*_*n*_ = [*U*_1,*n*_, …, *U*_*M,n*_]^T^ is a vector of *M* zero-mean white and uncorrelated input noise processes (prediction errors).

Starting from the time series of each given subject, the VAR model (1) was identified via a least squares method, finding estimates of the coefficients *A*_*k*_ and of the variance of the input noises, *i*.*e*.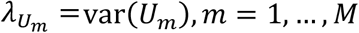, *M* (Faes et al., 2012). Afterwards, to compute Granger causality from the process *Y*_*i*_ to the process *Y*_*j*_, a so-called reduced model was defined and identified via the state-space method (Barnett and Seth, 2015), where the present state of the assigned target *Y*_*j,n*_ is described from the past states of all processes except the assigned driver *Y*_*i*_. This method avoids problems related to the incomplete formulation of the reduced model that may lead to poor estimates of GC (Faes et al., 2017). Denoting the prediction error process of the reduced model as *W*_*j*_ and its variance as 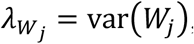, the GC correlation measure from *Y*_*i*_ to *Y*_*j*_, conditioned on the remaining processes *Y*_*k*_, *k* = {1, …, *M*}/{*i, j*}, is computed (Geweke, 1984):

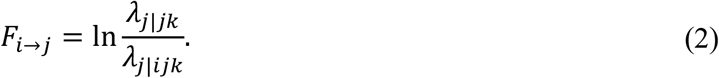

This conditional GC measure (2) quantifies the extent to which the past states of the driver *Y*_*i*_ help in predicting the present state of the target *Y*_*j*_ above and beyond its own past states and the past states of the other processes *Y*_*k*_. Here, the measure was computed by varying *i* and *j* in the range {1, …, *M*} (*i* ≠ *j*) to quantify the causal coupling along the two directions of interaction for each pair of glomeruli in each subject. Furthermore, to establish the existence of a link from the *i*^*th*^ to the *j*^*th*^ node of the observed network, the statistical significance of the computed conditional GC was tested using an asymptotic statistics approach. This approach makes use of a Fisher *F*-test applied to the estimated prediction error variances of the full and restricted models, considering the computation of multiple GC via false discovery rate (FDR) correction (α = 0.05) (Hochberg and Tamhane, 1987).

### Network analysis

For the analysis of signal dynamics across time, as a first step, the time series were sub-divided into seven 1-second windows: ON phase (0-1 s), earlyOFF phase (1-2 s), and OFF phases (2-3 s, 3-4 s, 4-5s, 5-6s, and 6-7s). Then, the signals belonging to the different windows were decomposed into slow and fast components by using an infinite impulse response (IIR) autoregressive filter with zero phase (Nollo et al., 2000) applied to the 30 trials. The filter was implemented first in a low-pass configuration (cut-off frequency ∼ 1.5 Hz) to extract the slow signal components mostly related to the stimulus response, and then in a high-pass configuration (cut-off frequency ∼ 2.5 Hz) to extract the fast components. In the latter case, before high-pass filtering, the mean response obtained by averaging the signal across the 30 repetitions was subtracted from each trial. As a last step, Granger causality analysis was performed for each bee, time window, and odorant to assess the causal connectivity structure of each network. The obtained matrices contain statistically significant GC correlation measures between glomeruli, or zero if results were not non-significant. The density of each estimated weighted network was evaluated as the fraction of significant connections to the total number of possible connections (Rubinov and Sporns, 2010). Significant changes in network density across time bins were assessed with a Kruskal-Wallis test performed separately for the unfiltered signal, the fast and the slow components.

### Stereotypy analysis

The similarity between the edge-centered odor representations within and across individuals was quantified via a best-match-to-template analysis. For the within-subject analysis, we split the 30 olfactory stimulations into 6 groups of 5 trials. Hence, GC measures were calculated independently for each group to obtain 6 connectivity maps for each odorant-bee combination. Then, the connectivity map for each stimulus trial (test) was tested against the mean response map across all remaining trials for all odorants (templates). The similarity between maps of two odors *o*_1_ and *o*_2_ was quantified by Pearson’s correlation coefficient *r* between the vectorized GC maps of odors *o*_1_ and *o*_2,_ after statistical validation. The best-fitting template is considered the one with the highest *r*. The probability of the best fit to the correct odor template is plotted in **Figure 2A**.

For the across-subject analysis, the average connectivity in an odorant-bee combination (calculated across all 30 olfactory stimulation) was tested against the average connectivity maps across all individuals excluding the tested one. For each test, the best template odorant would receive a score of 1 and all other odorants a score of 0. If the highest correlation was shared by more than one odor template, the score was split accordingly. Results in each bee and the averages across bees are shown in **Figure 2C**.

The same best-match-to-template test was performed for the node-centered maps, that is the time-averaged glomerular response amplitudes, within (**Figure 2B**) and across (**Figure 2D**) individuals.

For each test, significant differences from a random distribution, which would give 1/6 = 0.167 fit-to-template probability for each odor, were analyzed with a Wilcoxon sum rank test with FDR correction. Original and adjusted *p*-values for all tests and the signed rank values *W* are shown in **Supplementary Table 1**.

## Results

### Node-centered odor response maps

Glomerular activity was imaged by fast two-photon microscopy (Paoli et al., 2017) from projection neurons labeled with the calcium-sensor Fura-2. Spatio-temporal activity patterns were recorded as relative fluorescence changes, with respect to the baseline, upon stimulation with six odorants (*n* = 15 bees). Glomeruli were identified across animals by comparing their morphology and anatomical location with the honey bee AL reference atlas (Galizia et al., 1999a), and their odorant specificity (Galizia et al., 1999b). **Figure 1A** shows the trial-averaged response maps of the ten most frequently identified glomeruli across all bees, revealing their highly stereotyped response patterns. On average, two to four of the monitored glomeruli showed a clear excitatory response during odor arrival. This analytical approach for studying olfactory coding takes into account the responses of individual functional units, measuring physiological parameters such as glomerular response intensity or latency. It is the classical description of a stimulus representation in the AL: a node-centered network analysis.

**Figure 1.**
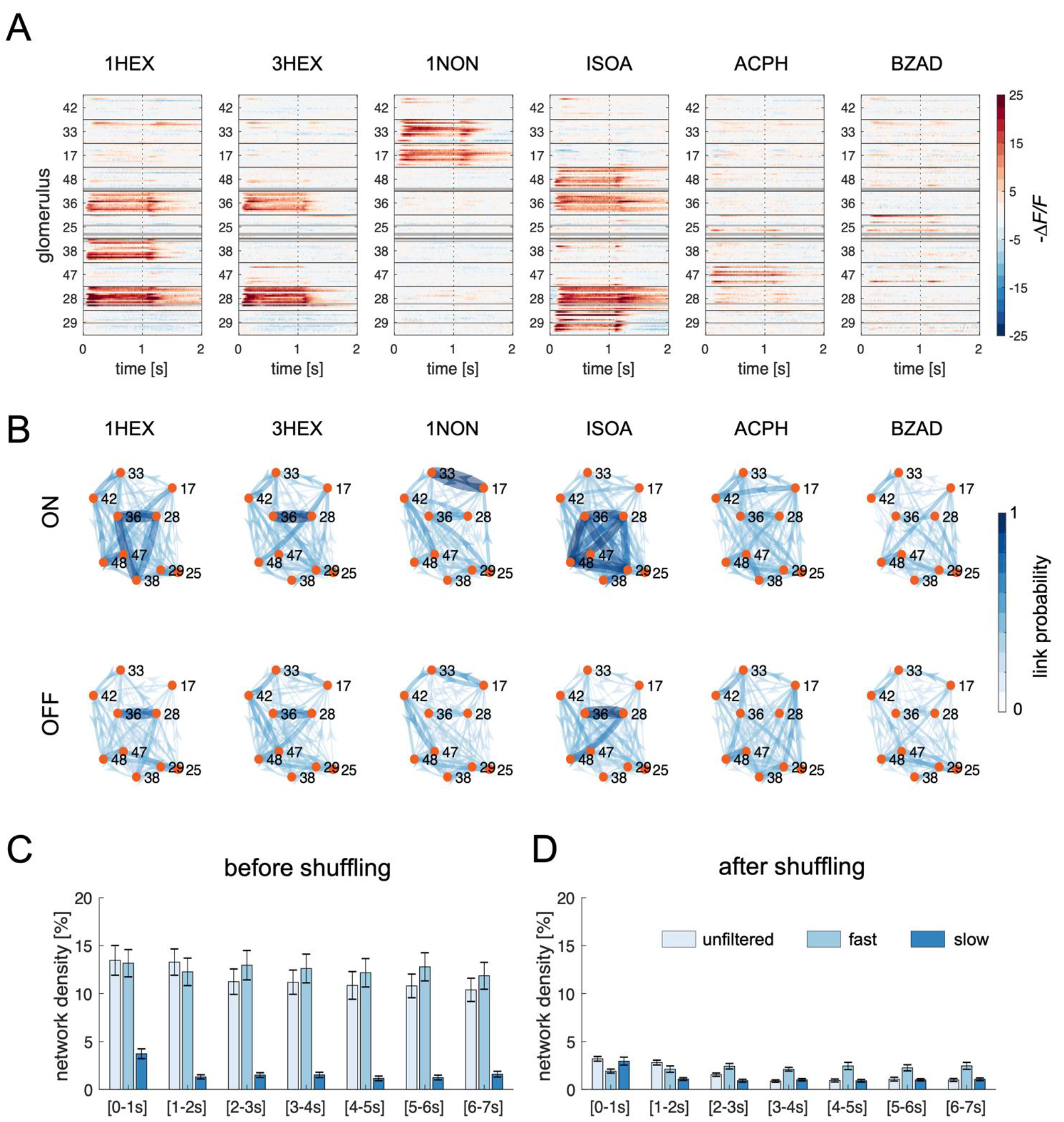
Node- and edge-centered odorant response maps. (**A**) Glomerular responses across bees and glomeruli. The relative fluorescence change across time is color-coded; gray lines represent the unavailability of individual glomerular data in single bees. Olfactory stimulation is delivered in the 0 to 1 s interval. The *y*-axis shows the response profiles of individual bees (*n* = 15) grouped according to the glomerulus ID number. (**B**) Mean connectivity maps across all bees calculated during stimulation (*t* = 0 to 1 s, top row) and 5 s after odor offset (*t* = 6 to 7 s, bottom row). Link thickness and color darkness indicate the probability of link detection across all analyzed bees. (**C**) Mean network density averaged over all bees, odorants, and time for connectivity maps computed from the unfiltered signal, the fast, and the slow signal components. (**D**) Network density of the connectivity maps calculated from the same dataset after shuffling glomeruli across individuals. Abbreviations: 1-hexanol, 1HEX; 3-hexanol, 3HEX; 1-nonanol, 1NON; isoamyl acetate, ISOA; acetophenone, ACPH; benzaldehyde, BZAD.

### Edge-centered odor response maps

#### The nature and dynamics of inter-glomerular connectivity

High-temporal resolution allows investigating the temporal correlations between glomerular responses, as shown here for the first time. This provides access to the functional connectivity across glomeruli, shifting the focus from the network nodes, the AL glomeruli, to its edges, the inter-glomerular connections. The connectivity analysis was performed via Granger causality, in its conditional form, which quantifies the amount of information contained in the present state of one glomerulus (target process) that can be predicted by the past state of another glomerulus (source process), above and beyond the information that is predicted already by the past states of the target process itself and of all others glomeruli (Barnett et al., 2009).

To investigate whether inter-glomerular connections are permanently active links within the AL, or if they are elicited by an olfactory input, we compared causal connectivity maps during and after olfactory stimulation. Note that with connectivity or edge-centered maps we refer to the calculated GC correlation matrices after statistical validation (see **Materials and Methods**). **Figure 1B** shows the GC correlation among the analyzed glomeruli during olfactory stimulation (ON, 0-1 s window) and 5 s after stimulus offset (OFF, 6-7 s window). Causal connectivity maps represent the average inter-glomerular connectivity detected across all measured honey bees, and connections (color and thickness-coded) are normalized from 0 (no individual shows that connection) to 1 (all animals present that specific inter-glomerular connection). As shown, edge-centered maps are rich in connectivity both in the presence and absence of olfactory input. Nevertheless, the network density decreases significantly after odor offset, suggesting that olfactory stimulation modulates inter-glomerular connections (**Figure 1B** and **C**).

A comparison between node- and edge-centered maps revealed that causal connections are more likely to link strongly responsive glomeruli. To test whether GC depends on glomerular response patterns similarity, rather than on information transfer across nodes, we used complementary approaches:

First, we filtered glomerular activity traces in slow and fast components (see **Materials and Methods**): the former is dominated by slow stimulus-induced transients, while in the latter these components are removed so that it consists only of fast oscillations (**Supplementary Figure 1**). If network states were dominated by the similarity between glomerular responses, the links between highly responsive glomeruli should be present also in the slow signal, deprived of fast oscillatory components. However, GC connectivity analysis of the slow components provided a reduced number of edges, advocating against an equivalence of node- and edge-centered odor representations (**Figure 1C**). Conversely, analyzing only the fast components, the network density remains comparable to the one in the unfiltered signal, suggesting that fast oscillations have a dominant role in providing inter-glomerular connectivity information.

Then, we tested whether the identified connections between glomeruli were dependent on response similarity, thus providing redundant information to the node-centered odorant representation (**Figure 1A**). Thanks to the highly conserved glomerular response pattern across bees, shuffling glomeruli among individuals provides artificial AL networks with similar glomerular response profiles. Interestingly, a GC analysis of the shuffled data generated connectivity maps with a dramatic decrease in network density across bees and stimuli (**Figure 1D**), demonstrating that similarity between glomerular response profiles is not sufficient to produce the observed causal functional connectivity. Finally, we investigated connectivity map dynamics by measuring changes in connectivity during and after odor stimulation. As illustrated in **Figure 1C**, the average network density remains high also after odor offset, although revealing a significant decrease 1 s after stimulus termination (Kruskal-Wallis test, unfiltered signal, trial effect *p* = 0.002). A similar trend affects the already low network density calculated from the slow components (*p* < 0.001), while no change could be detected in the fast component network density (*p* = 0.89).

Together this shows that measured causal functional connectivity is not representing response map similarity and slow transients, but physiological information transfer across glomeruli dominated by fast oscillatory components.

#### Causal functional connectivity contains odor information

The decrease in connection density over time supports the assumption that the observed inter-glomerular connections may be stimulus-induced and may provide stimulus-related information. To assess if information transfer across glomeruli is indeed stimulus-specific, we performed a series of best-match-to-template tests between GC matrices (see **Materials and Methods**) to determine whether a causal connectivity map can predict odor identity within and across individuals.

First, exploiting the fact that each bee was stimulated 30 times with each odorant, we split each recording into six groups of five trials. Then, we performed the GC analysis on each group of trials, obtaining six independent connectivity maps for each bee/odorant combination. This allowed assessing how a GC connectivity map induced by a certain odorant is preserved within the same individual. **Figure 2A** shows the results for the best-match-to-template tests performed on connectivity maps calculated during olfactory stimulation (ON), immediately after (earlyOFF), and 5 s after odor offset (OFF). During olfactory stimulation, individual responses show a rate of odor recognition, which is well above chance level. This correct template identification is reduced right after odor offset - but stays significant for four odorants - and decays to chance level in the OFF window. For comparison, we assessed the probability of a correct template recognition using the node-centered glomerular response pattern (**Figure 2B**). As expected, stimulus responses within individuals are highly reproducible, resulting in a correct template recognition in the majority of the cases.

**Figure 2.**
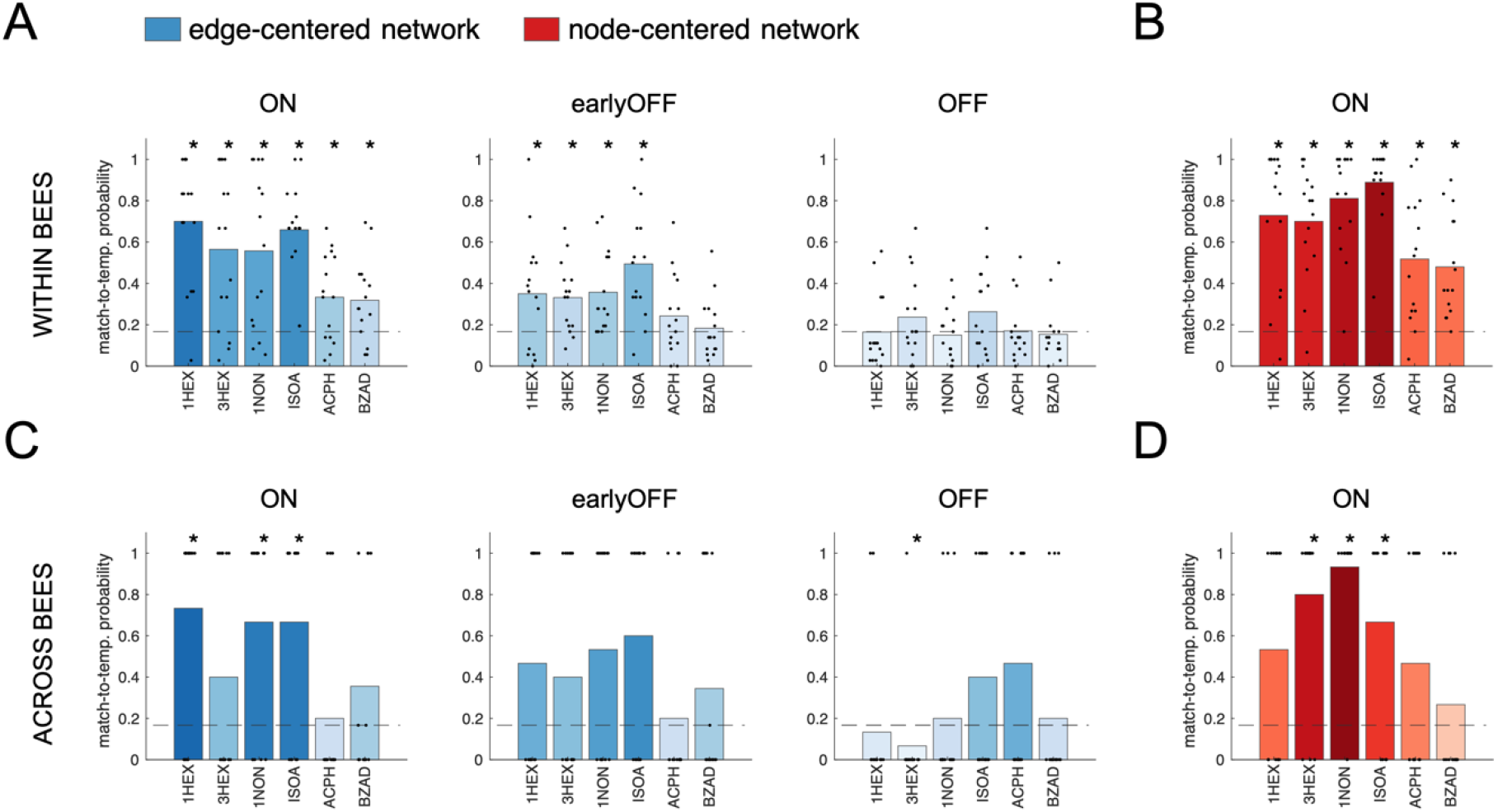
Best-match-to-template tests for edge-centered connectivity maps (**A** and **C**) and for node-centered glomerular response maps (**B** and **D**). (**A**) Within each bee, 6 connectivity maps were calculated for each odorant, each based on 5 stimulus repetitions. Each connectivity map (test) was tested against the mean map for each odor (templates), except the tested one. (**B**) Within each bee, the glomerular response map for each stimulus trial (test) was tested against the mean response map across all trials except the tested one for each odorant (templates). (**C**) Across bees, the trial-averaged connectivity profile for single odor/bee combinations (test) was tested against the mean connectivity profile of the individual odorants averaged across all bees except the tested one (templates). (**D**) Across bees, the trial-averaged glomerular response map for each odor-bee combination (test) was tested against the mean response maps of each odorant averaged across all bees except for the tested one (templates). In all subplots, dots indicate the test-to-template matching probability for each bee (*n* = 15); bars indicate the average value across bees. The horizontal dashed line represents the chance level. Significant differences from a random distribution were determined via a Wilcoxon signed rank test with FDR correction (* indicates adjusted *p*-values < 0.05, exact values in **Supplementary Table 1**).

Next, we investigated to what extent edge-centered maps are preserved across bees, *i*.*e*. if stimulus-related information exchange in the functional connectivity maps is similar at the population level. Hence, we calculated the individual connectivity map for each bee/odorant combination and tested if the connectivity maps of each bee for a test odorant could identify the correct subject-averaged odor template. Notably, across bees the best-match-to-template shows a higher variability than within animals, but is still significantly different from random matching for three stimuli (1-hexanol, 1-nonanol, and isoamyl acetate; **Figure 2C**). Again, we compared this result to the same test performed with the node-centered maps. **Figure 1A** suggests that the latter should be rather well-preserved across animals. Still, a match-to-template assay across individuals revealed that also in this case only three out of six odorants provide a response map that allowed identifying the correct template significantly better than chance (3-hexanol, 1-nonanol, and isoamyl acetate; **Figure 2D**).

## Discussion

### Differences in edge- and node-centered network representations

Since the observed connectivity is dominated by functional links among responsive glomeruli which are similar in all bees, we first verified whether it depends at all on physical links among glomeruli. By shuffling glomeruli across ALs, we generated a series of artificial ‘scrambled’ networks, which have node-centered maps similar to the original ones because of the stereotyped odor tuning of identified glomeruli across individuals. Granger causality analysis on scrambled AL provided a dramatically low number of connections, showing that glomerular response profile similarity is not sufficient to determine a causal functional connectivity link, and that dense connectivity maps could be obtained only among nodes belonging to the same physical network. In a previous report, Galan *et al*. showed that glomeruli displaying a strong odorant-induced response tend to fire more synchronously, possibly increasing their reciprocal excitatory connections (Galán et al., 2006). In agreement with the current observations on GC analysis, this advocates for a stimulus-induced functional connectivity and information transfer across responsive glomeruli.

Moreover, an analysis of the contribution of fast and slow signal components to the connectivity maps revealed that AL connectivity is dominated by fast oscillatory components, while the slow calcium transients - typical of slow signal components - have a minor role. Notably, the network density of the unfiltered signal is higher during olfactory stimulation but remains constitutively elevated also after stimulus offset. However, the odorant-related information content is confined to the stimulation phase, suggesting that even if active connections are always present, odorant arrival increases edge density in a stimulus-specific manner.

### Stimulus-dependence of edge- and node-centered network representations

Whether the observed connectivity maps contain stimulus-related information was assessed at the individual level by testing if the connectivity profiles calculated for the same bee but upon different olfactory stimulations could identify the correct odor template. Interestingly, repetition of the same stimulus provided odorant-specific connectivity networks, suggesting that information transfer among glomeruli - which determines the detected connectivity - carries stimulus-specific information. A similar approach was used to assess to what extent stimulus-elicited edge-centered maps are conserved across individuals. Despite the stereotypical responses of glomeruli across individuals, quantitative match-to-template tests did not allow for a general error-free odorant prediction for both node- and edge-centered maps. This is likely limited by the small sub-population of AL glomeruli we were able to identify with certainty. In fact, in a larger subset of glomeruli, each odorant would likely have some strongly responding units in its response map, possibly leading to a better match-to-template probability of the node-centered representation maps. Similarly, a larger subset of glomeruli could provide more stimulus-specific functional connections, resulting in more odorant-specific edge-centered maps. Moreover, the analysis was performed only on ten T1 glomeruli that could be faithfully identified across all tested bees, and it is possible that connectivity rules may be different across glomeruli from different tracts or when calculated on different sizes of glomerular populations. Altogether, within and across animal comparisons highlighted the presence of stimulus-related information in the functional connectivity maps, which are dominated by individuality. Nonetheless, template-identification tests performed across individuals based on node- and edge-centered maps were comparable. This suggests that connectivity maps are partly shaped by features that are conserved across individuals, thus allowing the recruitment of a similar set of inter-glomerular connections in response to the same odorant.

### Structure of the AL connectivity network

In a previous study on glomerular correlations, Linster and colleagues combined response amplitudes of AL input and output neurons and computational modeling to test different glomerular coupling scenarios. They found that neither a stochastic all-to-all coupling, nor spatially weighted connections favoring neighboring glomeruli matched the AL input/output data. Best matching to the experimental data was obtained with a functional model, where coupling was weighted by the similarity of the glomerular response profiles (Linster et al., 2005). Our measurements on temporal correlations via Granger causality analysis support these observations, showing that networks are not random, nor dominated by local connections, but provide stimulus specificity and include significant coupling of non-neighboring glomeruli. Moreover, the variability of stimulus-specific connectivity maps across subjects confirms previous physiological studies of local AL connectivity, which revealed that inhibitory connections varied across individuals, advocating that local connectivity is not pre-determined, but susceptible to interindividual variability that may depend on the difference in developmental plasticity and/or past experience (Girardin et al., 2012).

### Conclusive remarks

Olfactory input transformation within the AL relies on local connectivity, which plays a fundamental role in input gain control and contrast enhancement. Previous studies in honey bee and fruit fly revealed that the inter-glomerular connectivity is patchy (Girardin et al., 2012), and that local inhibitory connections participate in input signal gain control (Olsen and Wilson, 2008), while a network of lateral excitatory connections distributes odorant-evoked excitation between glomeruli (Olsen et al., 2007). Here, we showed that the application of a Granger causality approach to high-temporal resolution glomerular calcium imaging provides access to information on the functional connectivity among glomeruli, the network nodes, and allows monitoring how such connectivity changes during and after an olfactory stimulation. We demonstrated that GC-based connectivity describes physical coupling between neurons and observed that, although connections can often be found among strongly responsive glomeruli, response similarity is not sufficient to determine a functional connection between units. In the future, the use of pharmacology will be crucial in understanding the neurochemical nature behind such glomerular connectivity, and in particular the role of the local GABAergic interneurons (Schäfer and Bicker, 1986) in mediating between-nodes information transfer. and particularly the contribution of excitatory and inhibitory local neurons to the between-nodes information transfer. Similarly, extending the analysis to larger AL networks will provide further insight into the information content of edge-centered maps and will allow assessing the degree of information transfer between glomeruli innervated by different antennal nerve tracts.

## Author Contributions

M.P. and A.H. planned and designed the study, M.P. performed the experiments, M.P. and Y.A. analyzed the data, A.H. and L.F. provided funding, equipment, and codes, and all authors contributed to writing the manuscript.

## Acknowledgments

We acknowledge the assistance of Simone Monachino with data analysis.

## Supplementary Information

**Supplementary Table 1.** Statistical test on best-match-to-template tests. Wilcoxon signed rank test parameters: original *p*-values (black), FDR-adjusted *p*-values (red), and signed rank value *W* (blue). For each odor/bee combination, *n* = 15.

|  | Edge- centered |  |  |  |  |  | Node-centered |  |
| --- | --- | --- | --- | --- | --- | --- | --- | --- |
|  | within |  |  | across |  |  | within | across |
|  | ON | earlyOFF | OFF | ON | earlyOFF | OFF | ON | ON |
| 1HEX | 0.0001 | 0.013 | 0.72 | 0.0016 | 0.16 | 0.078 | 0.0002 | 0.07 |
|  | 0.0004 | 0.020 | 0.72 | 0.0010 | 0.32 | 0.24 | 0.0002 | 0.11 |
|  | 119 | 103 | 52 | 110 | 84 | 29 | 118 | 92 |
| 3HEX | 0.0016 | 0.0022 | 0.30 | 0.45 | 0.45 | 0.0065 | 0.0001 | 0.0005 |
|  | 0.0032 | 0.0044 | 0.59 | 0.54 | 0.54 | 0.04 | 0.0002 | 0.0015 |
|  | 99 | 98 | 53 | 75 | 75 | 15 | 119 | 114 |
| 1NON | 0.0034 | 0.0005 | 0.54 | 0.0056 | 0.07 | 0.32 | 0.0001 | 0.0001 |
|  | 0.0050 | 0.0015 | 0.64 | 0.012 | 0.21 | 0.38 | 0.0002 | 0.0007 |
|  | 109 | 78 | 42 | 105 | 92 | 42 | 105 | 119 |
| ISOA | 0.0001 | 0.0004 | 0.12 | 0.0056 | 0.016 | 0.45 | 0.0001 | 0.0056 |
|  | 0.0004 | 0.0015 | 0.59 | 0.012 | 0.10 | 0.45 | 0.0002 | 0.012 |
|  | 120 | 103 | 78 | 105 | 99 | 75 | 120 | 105 |
| ACPH | 0.018 | 0.28 | 0.52 | 0.32 | 0.32 | 0.16 | 0.0010 | 0.16 |
|  | 0.018 | 0.33 | 0.64 | 0.47 | 0.47 | 0.32 | 0.0010 | 0.20 |
|  | 101 | 71 | 42 | 42 | 42 | 84 | 101 | 84 |
| BZDA | 0.011 | 0.99 | 0.22 | 0.56 | 0.68 | 0.32 | 0.0001 | 0.71 |
|  | 0.013 | 0.99 | 0.59 | 0.56 | 0.68 | 0.38 | 0.0002 | 0.71 |
|  | 92 | 61 | 27 | 55 | 60 | 42 | 105 | 54 |

**Supplementary Figure 1.**
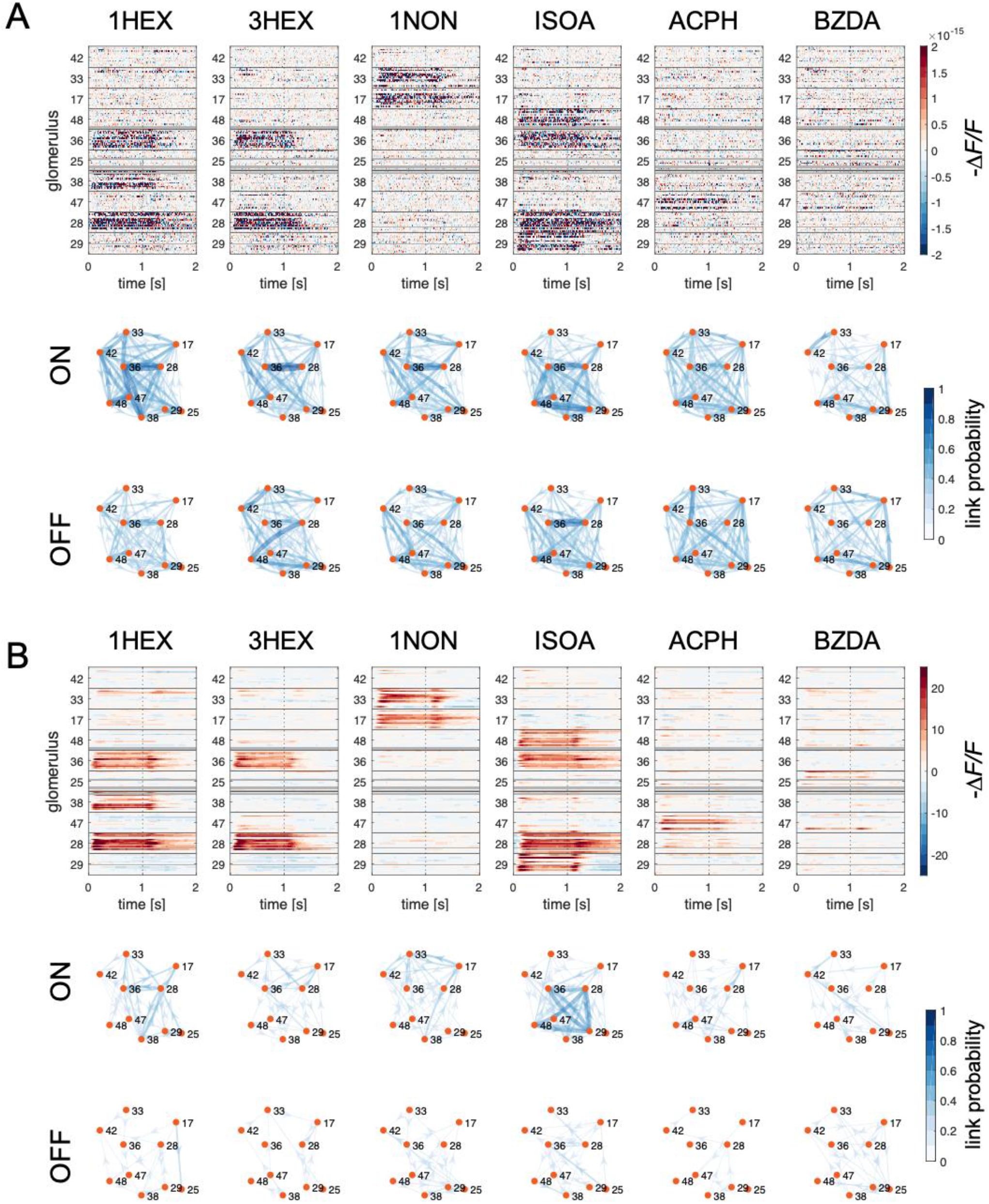
Glomerular profiles and connectivity maps computed from the fast (**A**) and slow (**B**) signal components. (**A**) Glomerular responses across bees and glomeruli. The relative fluorescence change is color-coded as a function of time, gray lines represent the unavailability of individual glomerular data in single bees. Olfactory stimulation is delivered in the 0-1s interval. The *y*-axis shows the response profiles of individual bees (*n* = 15) grouped according to glomerulus ID number. Mean connectivity maps across all bees calculated during stimulation (*t* = 0 to 1 s, top row) and 5s after odor offset (*t*=6 to 7s, bottom row). (**B**) The same data are presented after filtering original calcium signals to preserve only the slow components. Abbreviations: 1-hexanol, 1HEX; 3-hexanol, 3HEX; 1-nonanol, 1NON; isoamyl acetate, ISOA; acetophenone, ACPH; benzaldehyde, BZAD.

